# Plasma Proteomic Profiling Identifies a Subset of HIV Elite Controllers with Inflammatory Profiles Resembling People Without HIV

**DOI:** 10.64898/2026.09.06.749753

**Authors:** Shane D. Falcinelli, David P. Maison, Isaac Avila-Vargas, Thomas Dalhuisen, Chenyang Jiang, Xiaodong Lian, Ce Gao, Min-Gyoung Shin, Reuben Thomas, Jean Feng, James Salazar, Rebecca Hoh, Jeffrey N. Martin, Steven G. Deeks, Peter W. Hunt, Mathias Lichterfeld, Xu Yu, Timothy J. Henrich, Michael J. Peluso

## Abstract

**Background:** Elite controllers (ECs) are people with HIV (PWH) who suppress viral replication without antiretroviral therapy (ART). A subset, often termed exceptional elite controllers (EECs), harbor very small viral reservoirs, with few intact proviruses integrated into inactive chromosomal regions. Defining inflammatory signatures in ECs may clarify whether some approach a functionally cured state.

**Methods:** We measured 384 plasma proteins in ART-naïve ECs (n=62), ART-suppressed non-controllers (n=37), and people without HIV (PwoH) (n=50) using Olink proteomics, and compared proteomic profiles, clinical parameters, and intact reservoir data using FLIP-Seq. Hierarchical clustering identified ECs whose proteomic profiles most closely resembled PWoH; this subset was further analyzed using machine-learning approaches.

**Results:** ART-suppressed non-controllers had higher inflammatory protein levels than ART-naïve ECs, yet ECs still showed elevated inflammation and lower CD4:CD8 ratios compared with PWoH. Clustering analysis revealed a distinct subset of ECs (n=9) whose profiles closely matched those without HIV (ECs with PWoH-like proteomic profiles), a pattern not observed in ART-suppressed non-controllers. Compared to typical ECs, ECs with PWoH-like proteomic profiles had consistently lower IL-1β, SCRN1, and BID, numerically lower intact proviral frequencies, and a high frequency of protective HLA alleles. Machine-learning analyses highlighted features that most strongly differentiated ECs with PWoH-like proteomic profiles from typical ECs, including reduced gut mucosal/intestinal epithelial–associated proteins, higher absolute CD4 counts, and lower levels of pro-inflammatory cytokines and cellular immune activation.

**Conclusions:** These findings identify a subset of ECs with a PWoH-like inflammatory proteomic profile, accompanied by numerically lower intact HIV reservoirs and a high frequency of protective HLA alleles. Defining the mechanisms associated with this low-inflammatory state may provide insights into exceptional HIV control and inform strategies for durable HIV remission.

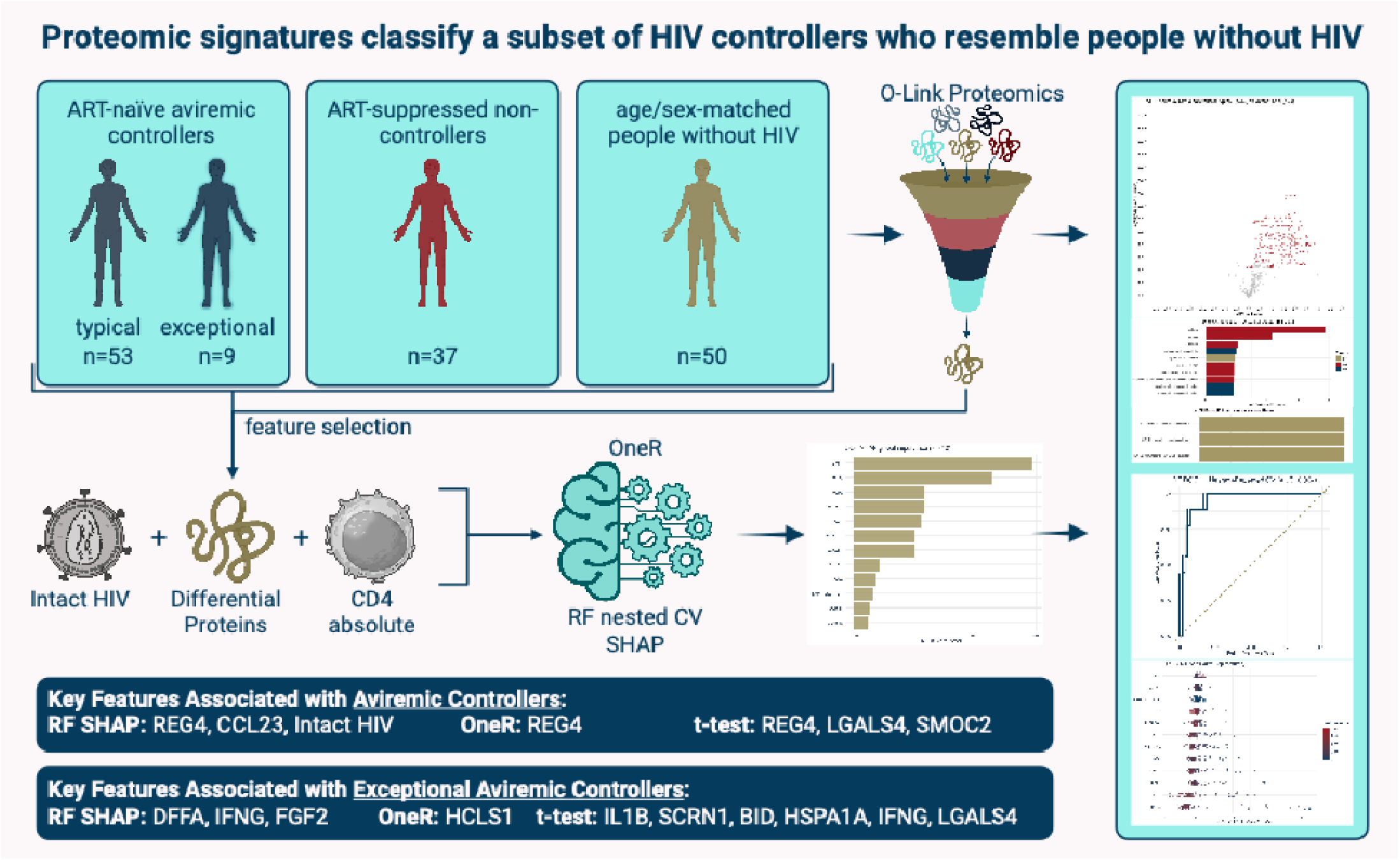

## INTRODUCTION

Understanding why a small subset of people with HIV can suppress viremia without antiretroviral therapy (ART) is a major focus of cure-oriented research [1]. In most people with HIV, untreated infection is characterized by ongoing viral replication and progressive immune dysfunction. In contrast, elite controllers (ECs) are typically defined by virologic criteria alone: they maintain plasma HIV RNA below the limit of detection of standard commercial assays in the absence of ART, allowing for occasional low-level blips [2, 3]. Although ECs are virologically defined, their immunologic trajectories and clinical outcomes can vary, and not all ECs maintain stable immune function over time [4–8].

Among ECs, clinical outcomes vary [8]. Some ECs have lower rates of non-AIDS morbidity and mortality than ART-suppressed non-controllers [4, 5]. However, other ECs experience gradual CD4⁺ T cell decline, and a subset eventually lose virologic control, often coinciding with impaired HIV-specific immune responses [6–9]. Even with sustained viral suppression, some ECs exhibit persistent immune activation and inflammation [10–12]. This may contribute to CD4⁺ T cell loss and improve with ART initiation [13, 14]. As a result, clinical guidelines recommend that ART be considered for ECs, particularly those with declining CD4 counts or evidence of loss of control of viremia [15]. These observations highlight the need for predictors of durable elite control beyond virologic criteria, including sustained immune health and low inflammatory burden.

More recently, an “exceptional” elite controller (EEC) phenotype has been described [16–23]. This phenotype is defined in some cases by decades-long ART-free suppression without clinical sequelae, a potential model for a functional cure [16–23]. Studies of exceptional elite controllers have described combinations of very small or undetectable replication-competent reservoirs, unusual proviral integration patterns including enrichment in transcriptionally inactive chromosomal regions, protective HLA class I alleles, and low levels of immune activation or inflammation [16–23]. However, key gaps remain in understanding the inflammatory profiles that distinguish ECs from non-controllers and determining whether there are distinct profiles within ECs. Defining these profiles may reveal therapeutic targets to reduce long-term inflammation in individuals without exceptional HIV control.

In this study, we used high-throughput plasma proteomics to characterize the inflammatory profiles of ART-naïve ECs, comparing them with ART-suppressed non-controllers and people without HIV (PWoH). We identified a subset of ECs whose plasma proteomic profiles closely resembled those of PWoH and termed them ECs with PWoH-like proteomic profiles. These findings provide new insights into the plasma proteomic landscape of HIV elite control and highlight candidate pathways associated with a PWoH-like inflammatory state that may inform future studies of HIV remission and cure.

## RESULTS

### Clinical and reservoir characteristics

The cohort characteristics are summarized in **Table 1**. Demographic, clinical, and proviral parameters were compared across groups [19]. Gender and CMV status were similar between ECs and ART-suppressed non-controllers. Median age was modestly higher in both groups of PWH than in PWoH (ECs +6 years; ART +8 years).

**Table 1.**
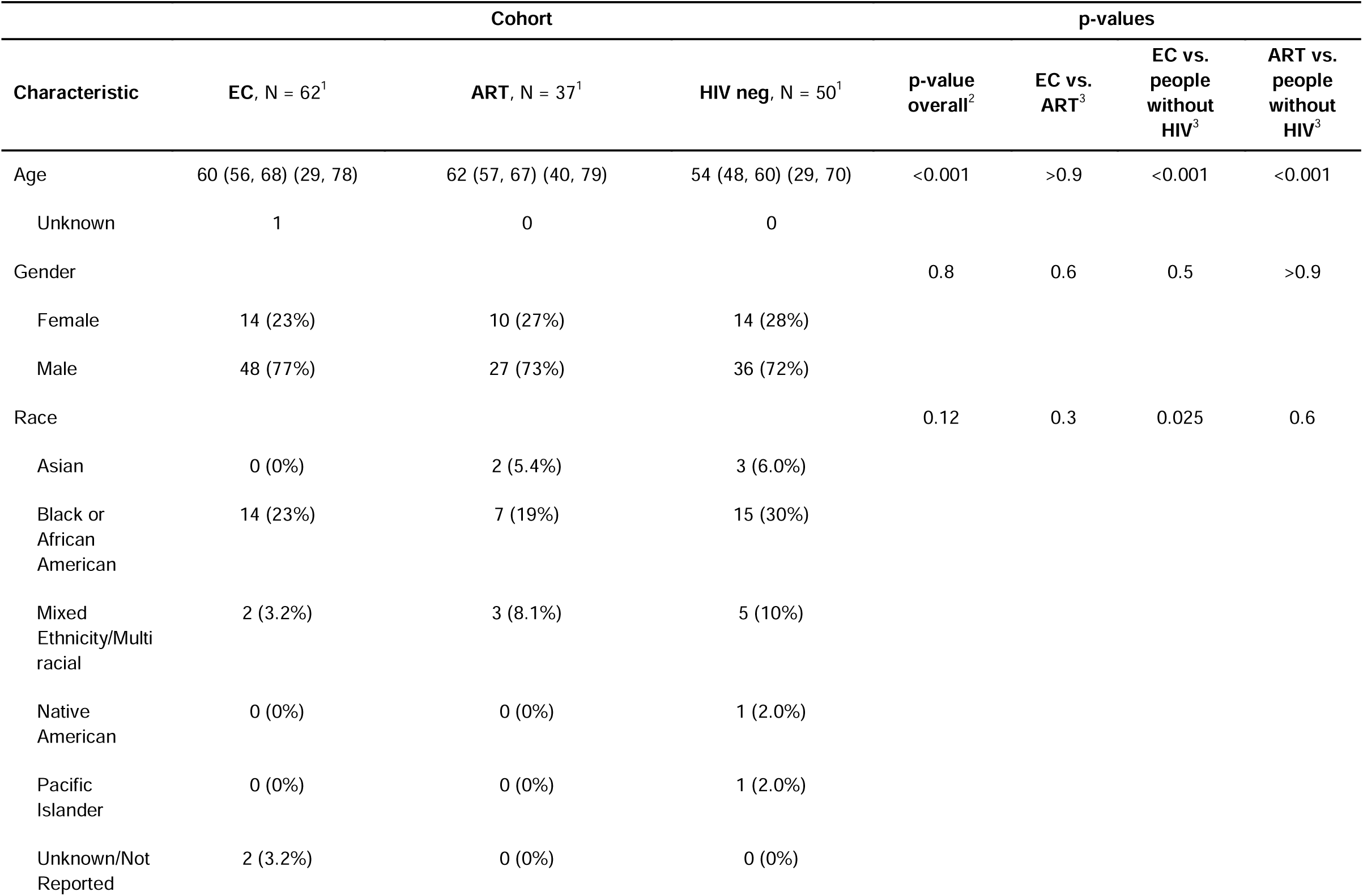
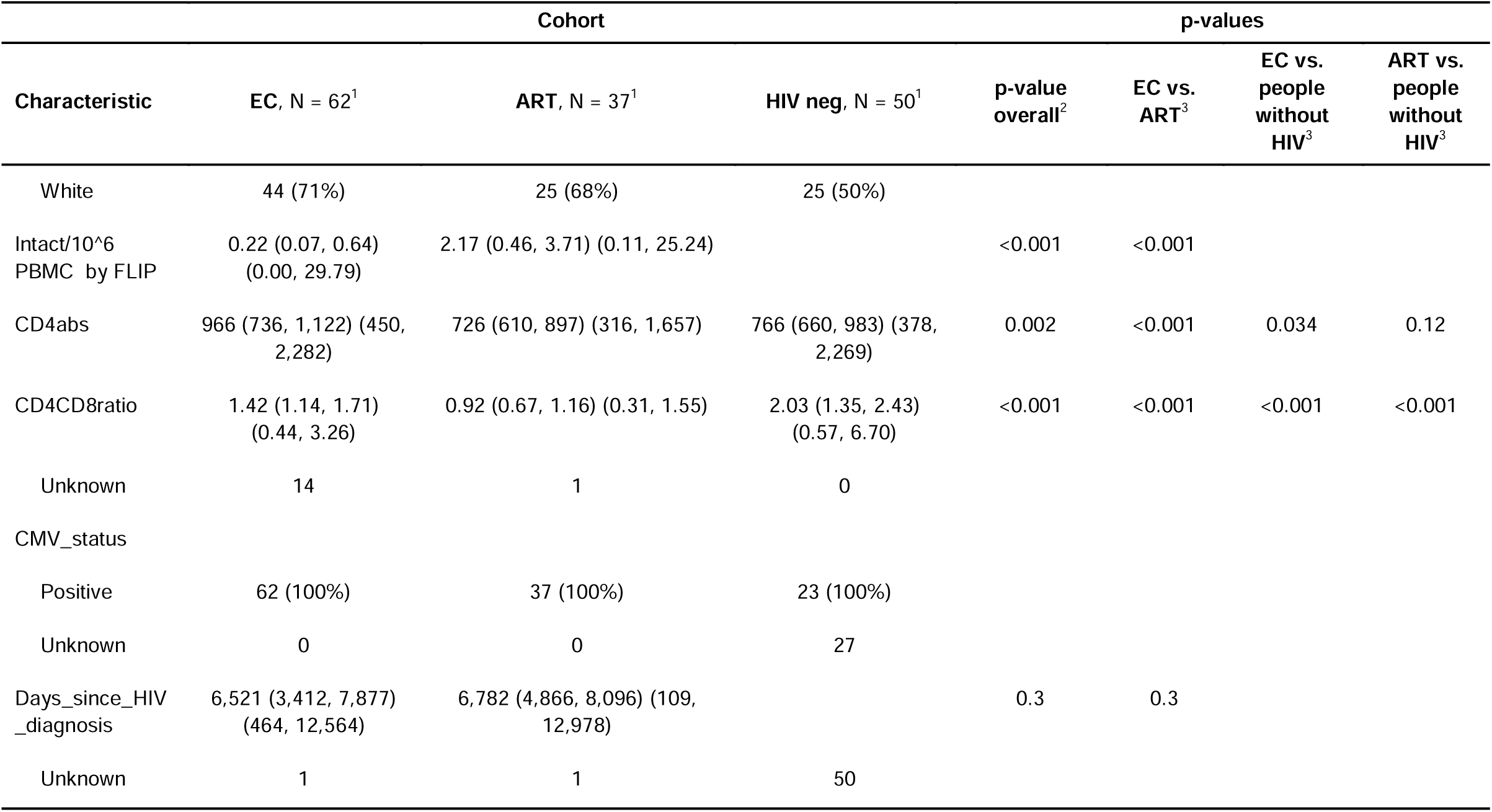
Demographics and clinical parameters. ^1^Median (IQR) (Range); n (%); ^2^Kruskal-Wallis rank sum test; Pearson’s Chi-squared test; Fisher’s exact test; ^3^Wilcoxon rank sum test; Fisher’s exact test.

As previously reported, ECs had significantly lower intact HIV DNA frequencies by FLIP-seq than ART-suppressed non-controllers (p < 0.001; **Figure 1A**) [19]. ECs also exhibited higher CD4+ T-cell counts (p < 0.001; **Figure 1B**) and higher CD4:CD8 ratios (p < 0.001; **Figure 1C**). However, the groups had a similar duration since HIV diagnosis (**Figure 1D**). All results were consistent in unadjusted and age- and gender-adjusted analyses.

**Figure 1.**
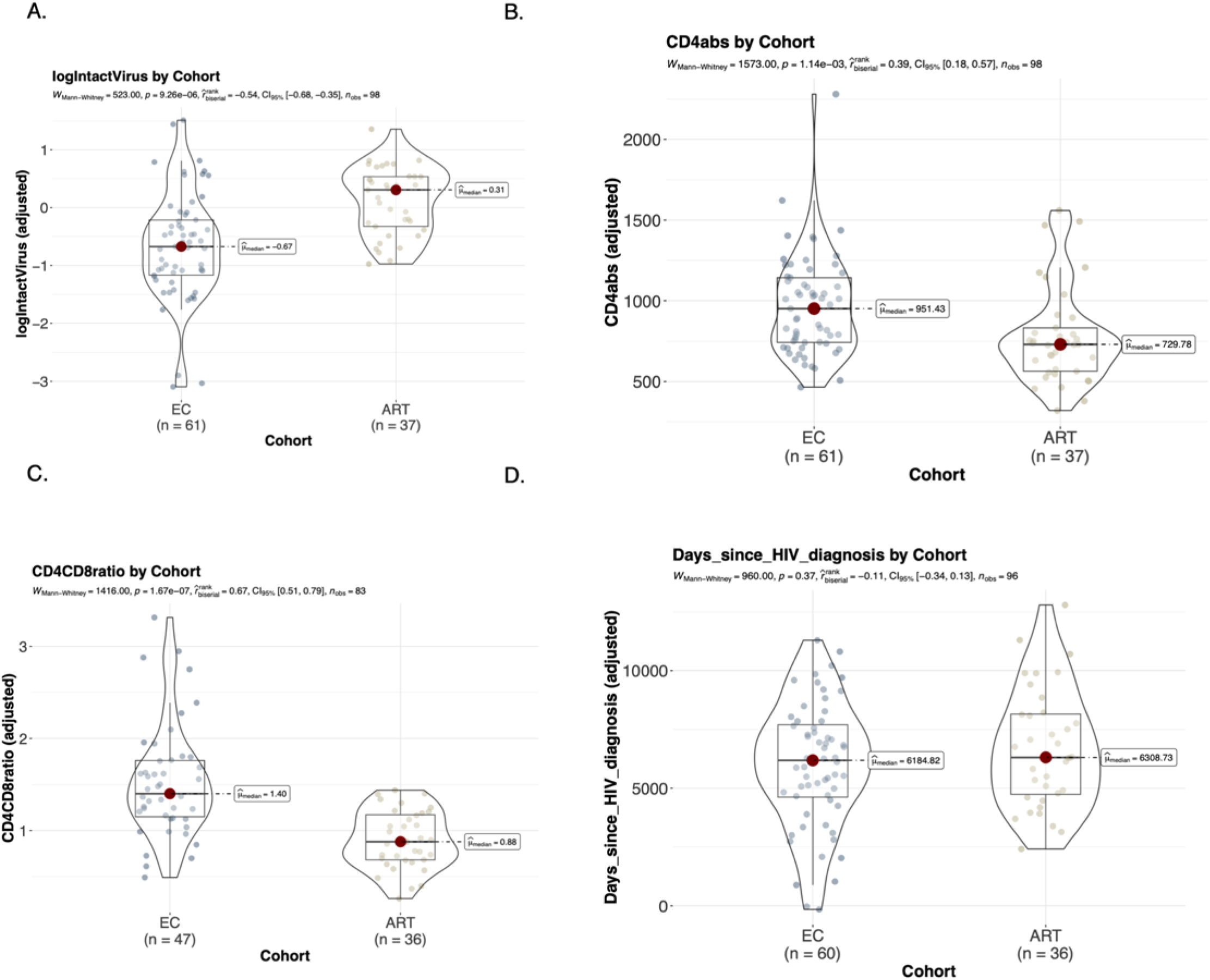
ART-naïve aviremic elite controllers (ECs) demonstrate higher CD4 counts, higher CD4:CD8 ratios, and lower intact proviral frequencies compared to ART-suppressed non-controllers. **A.** Mann– Whitney U test of intact HIV DNA frequencies shows significantly lower levels in ECs than in ART-suppressed non-controllers (p = 0.001). **B.** Mann–Whitney U test of CD4 counts shows significantly higher levels in ECs (p < 0.001). **C.** Mann–Whitney U test of CD4/CD8 ratios shows significantly higher ratios in ECs (p < 0.001). **D.** t-test of days since HIV diagnosis shows no significant difference between groups.

### ART-suppressed non-controllers have higher levels of inflammatory markers than ART-naïve ECs

We next compared plasma proteomic profiles between ART-suppressed non-controllers relative to ECs. We found 51 significant differentially expressed proteins between the two groups; 50 of 51 were upregulated in ART-suppressed non-controllers relative to ECs (**Figure 2A**). Qualitative analysis of the top upregulated proteins demonstrated increased chemokines (CCL23, CCL24, CCL25, CXCL12), extracellular matrix remodeling (SMOC2, AGRN, PRELP, CHRDL1), gastrointestinal epithelium-related proteins (REG4, LGALS4, SPINK4), and metabolism- and glycosylation-related proteins (CKMT1A/CKMT1B, B4GALT1). GO term enrichment analysis was used to identify biological processes and pathways overrepresented among differentially expressed proteins. The GO terms extracellular matrix, extracellular region, and structural molecule activity were enriched in ART-suppressed non-controllers relative to ECs **(Figure 2B).** Collectively, these findings highlight an inflammatory, extracellular matrix–remodeling protein signature that is enriched in ART-suppressed non-controllers relative to ECs.

**Figure 2.**
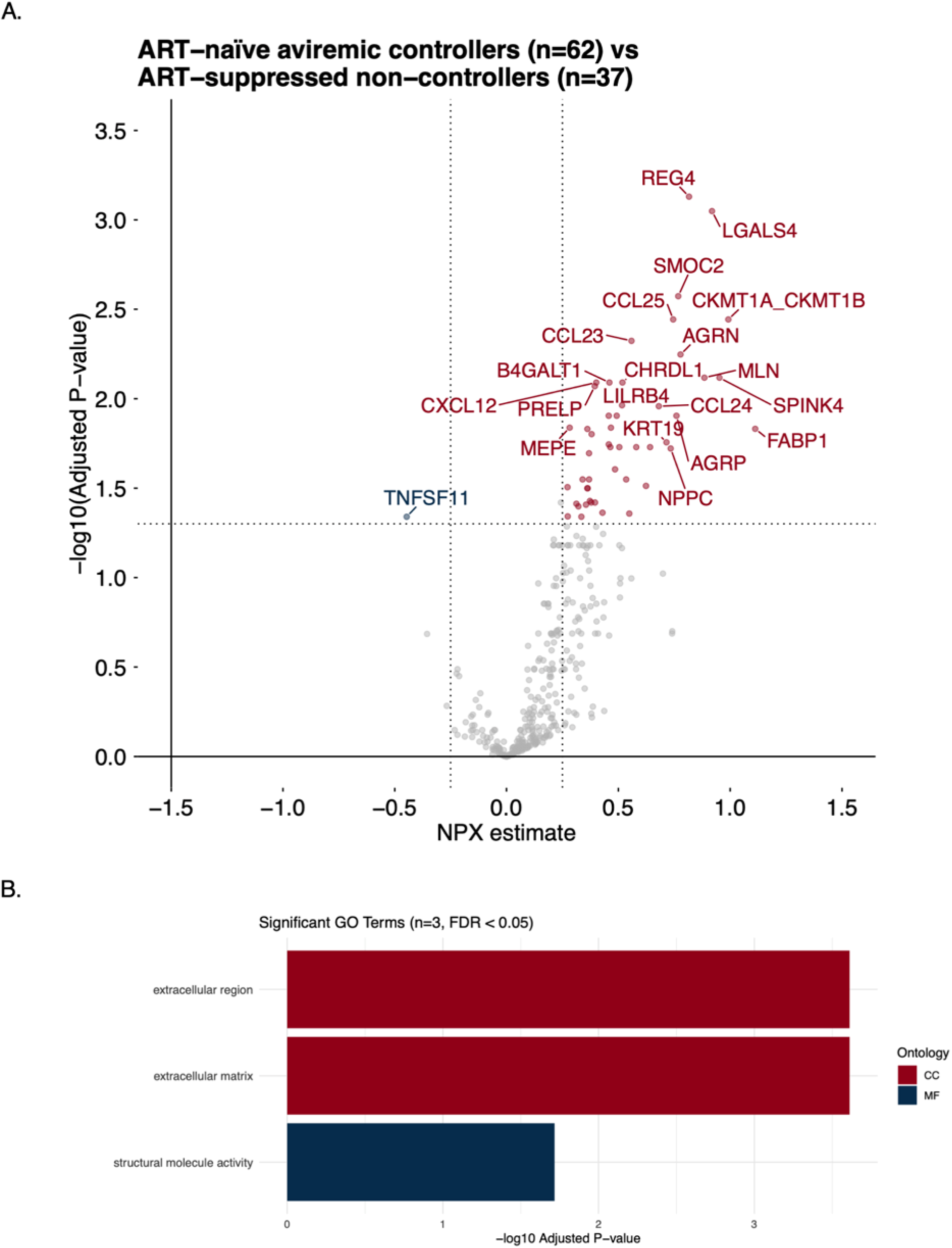
ART-suppressed non-controllers have higher levels of inflammatory markers than ART-naïve Elite Controllers (ECs). **A.** Volcano plot showing 51 differentially expressed proteins (DEPs) when comparing ART-suppressed non-controllers (n = 37) to elite controllers (n = 62), with 50 upregulated in ART-suppressed non-controllers and one upregulated in elite controllers (TNFSF11). The maximum estimate was 1.11, and the maximum –log10(adjusted p-value) was 3.13**. B.** Significant Gene Ontology (GO) terms enriched among the 51 DEPs.

### Elite controllers, as a group, have higher levels of inflammatory markers than PWoH

A parallel analysis compared plasma proteins in ECs and PWoH (**Figure 3**). Of 113 differentially expressed proteins, 110 were upregulated and 3 were downregulated in ECs relative to PWoH (**Figure 3A**). These proteins mapped largely to DNA repair (PSIP1, NBN), immune signaling and cytotoxicity (LSP1, PIK3AP1, SH2D1A, IL16, GZMB, NCF2), metabolism and stress response (EGLN1, PKLR, FOXO1), and apoptosis, proteostasis, and nucleotide metabolism (DFFA, SERPINB8, PSMG3, DNPH1). KEGG analysis identified enrichment of immune and infection-related pathway annotations, including NOD-like receptor signaling, B-cell receptor signaling, and the HIV-1 infection pathway (**Figure 3B**).

**Figure 3.**
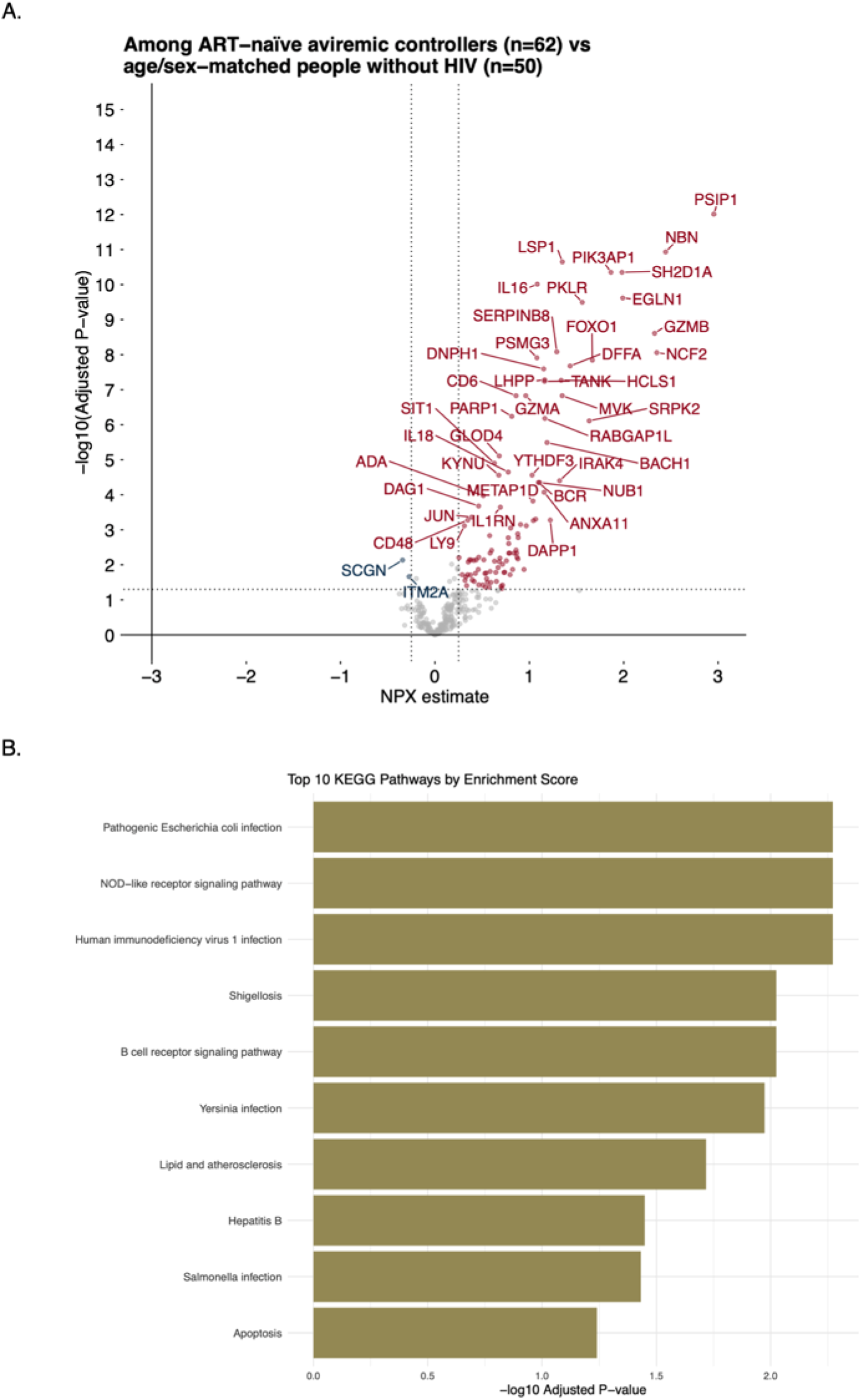
Elite controllers, as a group, have higher levels of inflammatory markers than people without HIV. **A.** Volcano plot showing 113 significantly differentially expressed proteins (DEPs) when comparing elite controllers (n = 62) to people without HIV (n = 50), with 110 upregulated in elite controllers and three upregulated in people without HIV (SCGN, ITM2A, and CD200). The maximum estimate was 2.94, and the maximum –log10(adjusted p-value) was 12.01. **B.** Top 10 Kyoto Encyclopedia of Genes and Genomes (KEGG) pathways ranked by adjusted p-values.

### Clustering analysis identifies a subset of ECs with similar plasma proteome profiles to people without HIV

Because ART-suppressed non-controllers showed higher inflammation than ECs (**Figure 2**), and ECs overall had higher inflammation than PWoH (**Figure 3**), we next asked whether a subset of ECs exhibited proteomic profiles similar to PWoH.

We performed three complementary clustering analyses of Olink NPX values to identify ECs co-clustering with the predominant PWoH group (**Figure 4**). Proteins were clustered based on similarity in expression patterns, with HOPACH used to select the optimal number of clusters, ensuring that the resulting groupings reflected robust patterns rather than arbitrary choices.

**Figure 4.**
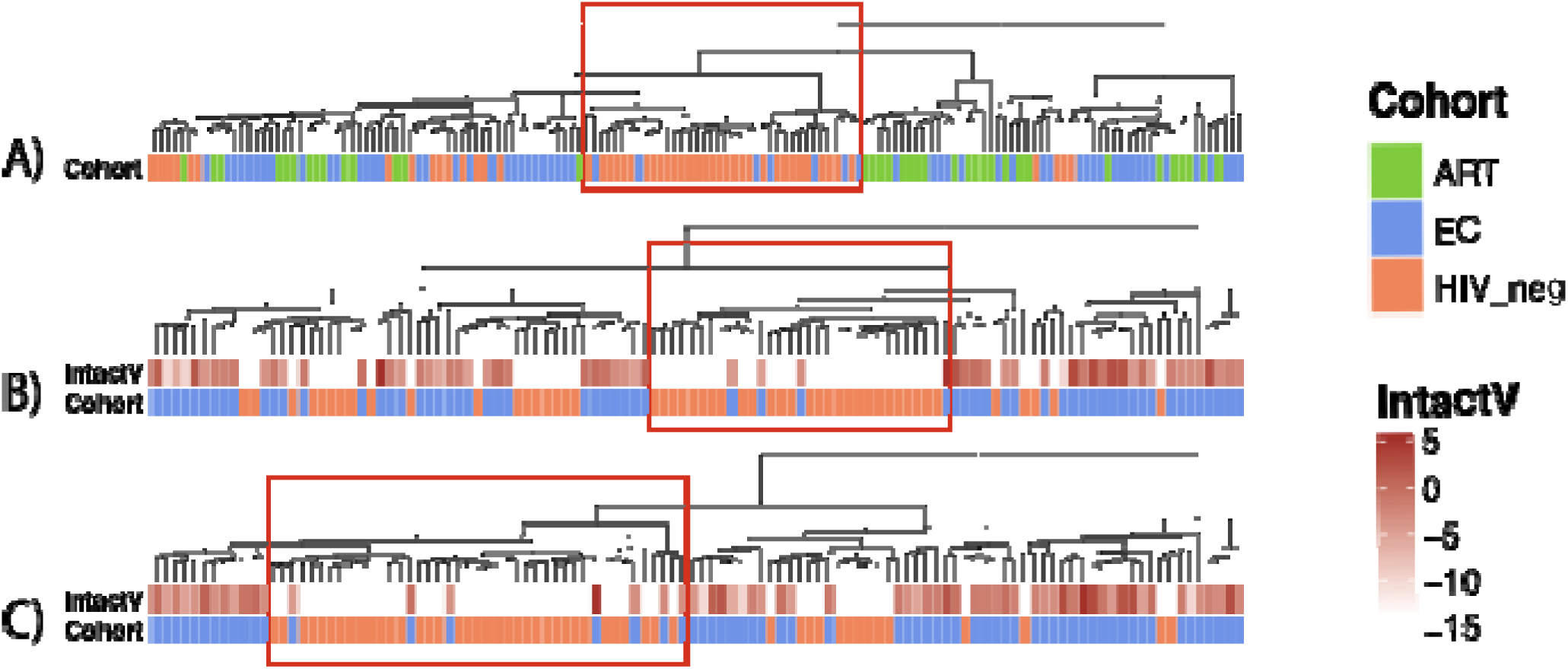
Clustering analysis identifies a subset of elite controllers (ECs) whose plasma proteomic profiles resemble those of people without HIV (PWoH). Three complementary clustering approaches were used to identify ECs with PWoH-like proteomic profiles, defined as ECs clustering within the largest HIV-negative group (orange). **A.** Clustering of ART-naïve aviremic ECs (n = 62, blue), ART-suppressed non-controllers (n = 37, green), and people without HIV (n = 50, orange) based on proteins differentially expressed by cohort after adjustment for gender. **B.** Clustering analysis repeated after removing ART-suppressed non-controllers, using all NPX values to compare ECs with HIV-negative individuals. **C.** A model-based clustering approach estimating NPX differences between ECs and HIV-negative individuals after adjusting for gender and age.

To ensure that we captured all PWH who might cluster with PWoH proteomically, we performed multiple clustering analyses (**Table 2**). First, clustering was performed using proteins differing by cohort after adjusting for gender (**Figure 4A**), identifying seven ECs that clustered with PWoH. We then repeated clustering using NPX values from ECs and PWoH. Four ECs co-clustered with PWoH using unadjusted data (**Figure 4B**) and seven did so after adjusting for age and gender (**Figure 4C**).

**Table 2.** ECs with PWoH-like proteomic profiles identified across three heatmap clustering analyses. Heatmap 1 represents clustering based on differential expression results for ECs with PWoH-like proteomic profiles identified through clustering using significant cohort-associated proteins derived from comparisons among elite controllers (ECs), ART-suppressed non-controllers, and people without HIV. Heatmap 2 corresponds to clustering based on differential expression results for ECs with PWoH-like proteomic profiles identified by unsupervised clustering with HIV-negative individuals using all measured proteins. Heatmap 3 corresponds to differential expression results for ECs with PWoH-like proteomic profiles identified by unsupervised clustering with HIV-negative individuals using all measured proteins, adjusted for age and gender.

| Participant Number | Heatmap 1<br>(Figure 4A) | Heatmap 2<br>(Figure 4B) | Heatmap 3<br>(Figure 4C) |
| --- | --- | --- | --- |
| 1 | Yes | No | Yes |
| 2 | Yes | No | Yes |
| 3 | Yes | Yes | Yes |
| 4 | Yes | Yes | Yes |
| 5 | No | No | Yes |
| 6 | Yes | No | Yes |
| 7 | Yes | Yes | Yes |
| 8 | Yes | No | No |
| 9 | No | Yes | No |

ART-suppressed non-controllers did not cluster with PWoH in any analysis (**Figure 4A-C**). Across all clustering approaches, nine ECs were identified as ECs with PWoH-like proteomic profiles (**Table 2**). We next assessed how these individuals differed from typical ECs in clinical parameters, reservoir frequency, and inflammatory signatures.

### ECs with PWoH-like proteomic profiles have numerically lower reservoir frequencies and frequently carry protective HLA alleles

We compared virologic and clinical parameters among the subset of nine ECs with PWoH-like proteomic profiles with typical ECs. There was a trend toward lower intact and total DNA HIV reservoir frequencies in ECs with PWoH-like proteomic profiles that clustered with people without HIV compared to those that did not (**Figure 5A-B**). ECs with PWoH-like proteomic profiles had a median intact proviral DNA frequency of 0.085/10^6^ PBMCs compared to 0.269/10^6^ PBMCs in typical ECs (Mann-Whitney Test, p = 0.24) and median total HIV DNA frequency of 0.692/10^6^ PBMCs compared to 1.622/10^6^ PBMCs in typical ECs (Mann-Whitney test, p = 0.25). No statistically significant differences or obvious trends in absolute CD4+ T cell count, CD4:CD8 ratio, or time since HIV diagnosis between the EC subsets were observed (**Figure 5C-E**).

**Figure 5.**
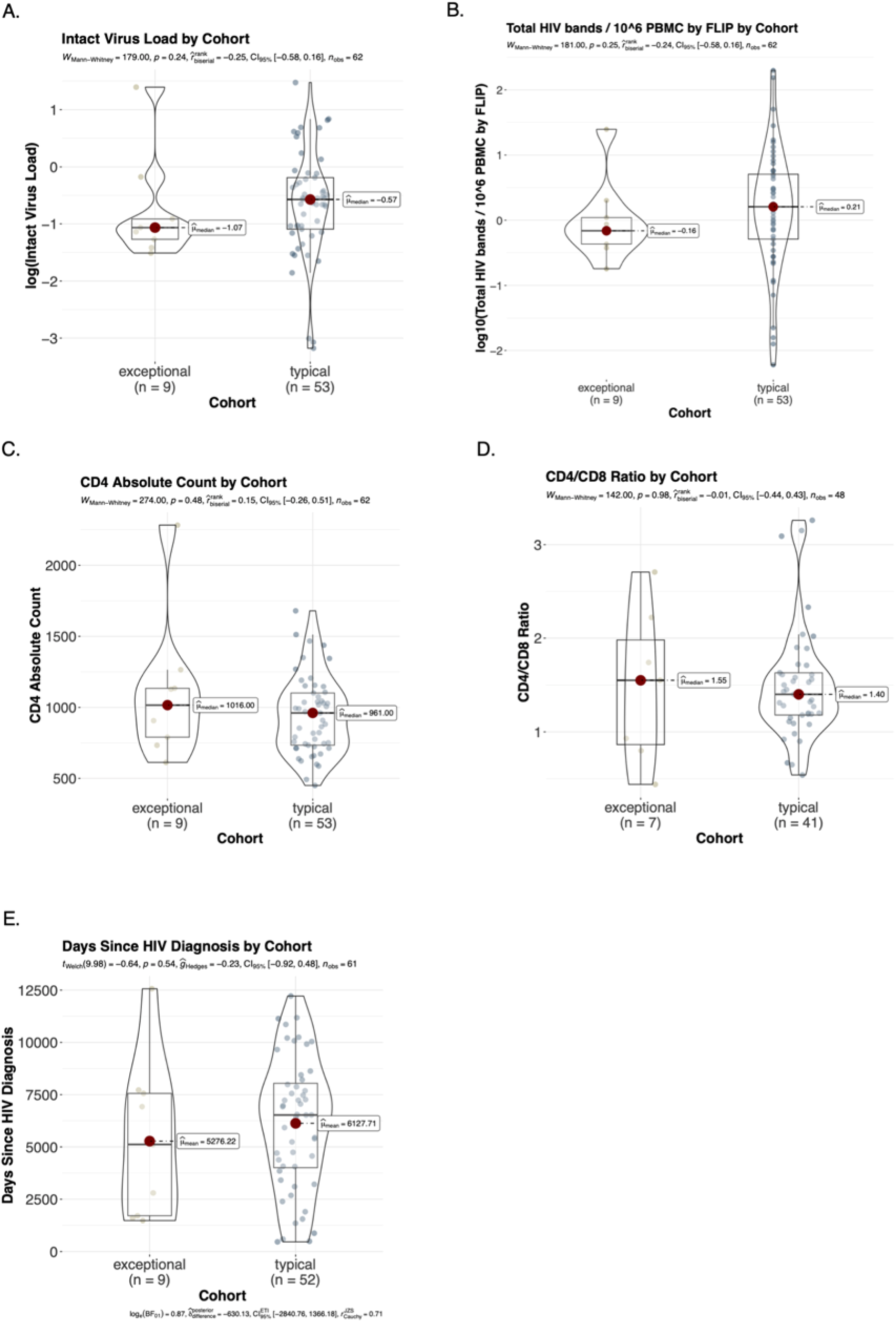
ECs with PWoH-like proteomic profiles tend to have lower intact and total HIV reservoir frequencies than typical ECs. **A.** A trend was observed toward lower intact proviral DNA frequency in ECs with PWoH-like proteomic profiles compared to typical ECs (Mann–Whitney test, p = 0.24). **B.** Total HIV DNA frequency was numerically lower in ECs with PWoH-like proteomic profiles than in typical ECs (Mann–Whitney U test, p = 0.25). **C.** No statistically significant differences in absolute CD4⁺ T cell count were observed between the subsets (p = 0.48). **D.** CD4/CD8 ratios did not differ significantly between groups (p = 0.98). **E.** Time since HIV diagnosis was also similar between the cohorts (Welch t-test, p = 0.54).

HLA class I genotyping showed that alleles previously associated with protection against HIV disease progression (HLA-B27:05, B57:01, B57:03 and B58:01) were more frequently observed among ECs with PWoH-like proteomic profiles [21, 24]. At least one protective allele was present in 7/9 ECs with PWoH-like proteomic profiles (78%), compared with 32/53 typical ECs (60%) and 8/37 ART-suppressed non-controllers (22%) (**Supplementary Table 1**).

### IL-1β, SCRN1, and BID are differentially expressed in ECs with PWoH-like proteomic profiles

To understand the protein features underlying the proteomically-defined group assignments, we examined which proteins were consistently differentially expressed between ECs with PWoH-like proteomic profiles and typical ECs across the different clustering analyses defined in **Figure 4A-C**. Because the ECs with PWoH-like proteomic profiles subgroup was defined by unsupervised analysis of the same Olink panel (**Figure 4A–C**), these comparisons are reported as characterization of the proteins distinguishing the predefined subgroups rather than as independent classification analyses

Qualitative analysis of the differential expression volcano plots identified additional proteins suggestive of increased inflammation and immunologic activation in proteomically-defined typical ECs compared to ECs with PWoH-like proteomic profiles (**Supplementary Figure 1**). These included classic pro-inflammatory cytokines (IL-12), markers of exhaustion (CD244), and stress-response/apoptosis (HSPA1A, PARP1, NBN, DFFA), all significant at adj. p < 0.05 in the comparison of proteomically-defined typical ECs compared to ECs with PWoH-like proteomic profiles. TNF was significant only in the age- and gender-adjusted subgroup (model 3, adj. p = 0.014).

Across all three clustering approaches (**Figure 4A-C**), IL-1β, SCRN1, and BID were among the top 15 most significantly differentially expressed proteins by adjusted p-value (**Supplementary Figure 1**). Higher expression by Olink NPX value of IL-1β, SCRN1, and BID was observed in typical ECs compared to ECs with PWoH-like proteomic profiles (**Figure 6A-C)**.

**Figure 6.**
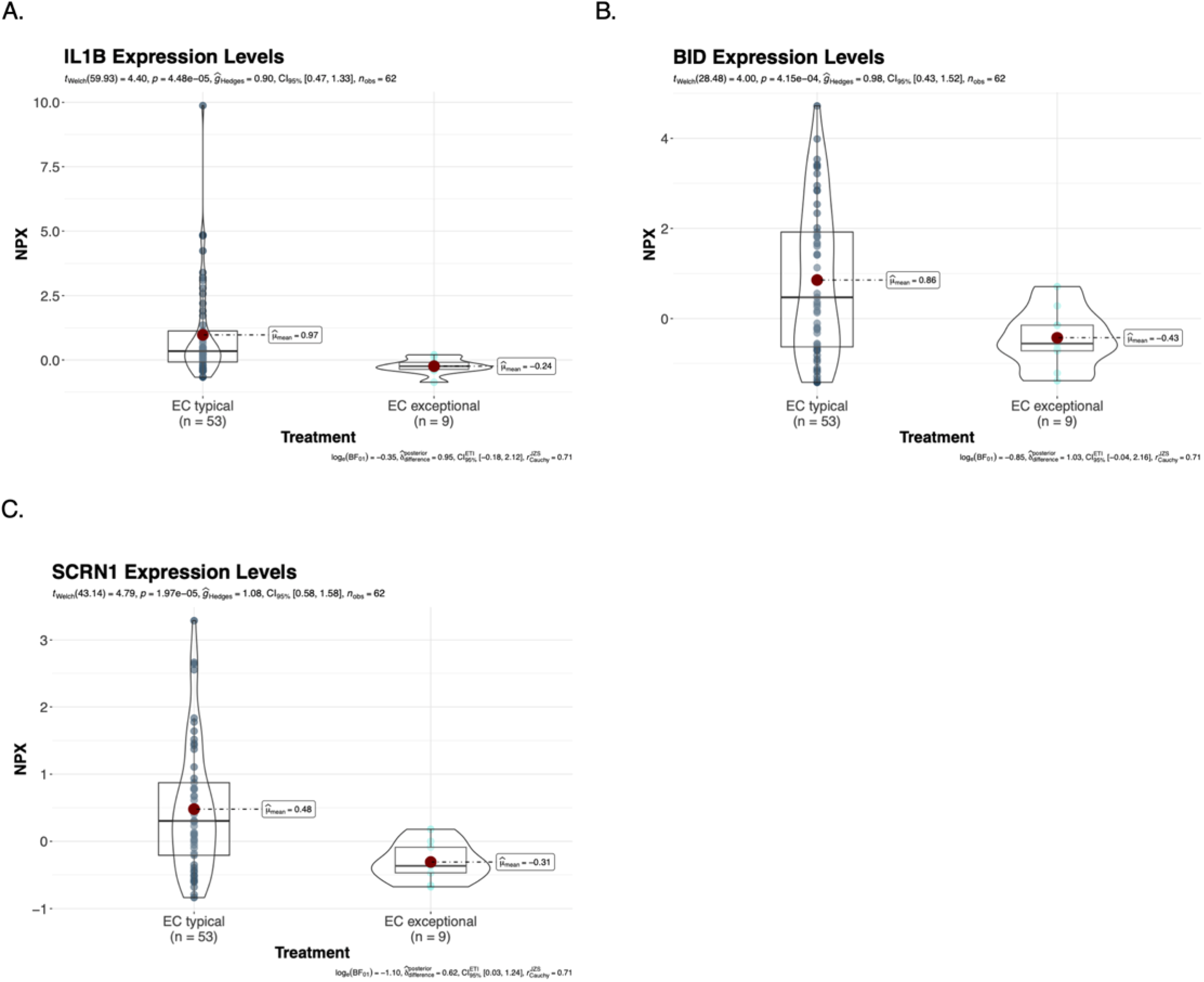
IL-1β, SCRN1, BID proteins are differentially expressed in ECs with PWoH-like proteomic profiles compared to typical ECs. To identify protein expression patterns that distinguish ECs with PWoH-like proteomic profiles from typical ECs, we compared results across three complementary clustering and differential expression approaches. IL-1β, SCRN1, and BID were consistently among the most differentially expressed proteins across the three clustering-defined comparisons, with all upregulated proteins enriched in typical ECs. Differential expression analysis comparing ECs with PWoH-like proteomic profiles (n = 9) versus typical ECs (n = 53) for **A.** IL-1β, **B.** BID, and **C.** SCRN1.

### Machine-learning analysis characterizes features distinguishing ECs from ART-suppressed non-controllers

To complement our differential expression analyses, we used a supervised machine-learning approach to characterize the combination of clinical and proteomic features that most strongly distinguished ECs from ART-suppressed non-controllers. Model inputs included Olink proteomic measurements, absolute CD4 counts, and intact proviral DNA. We used nested cross-validation Random Forest models with SHapley Additive exPlanations (SHAP) to assess the relative contribution of individual features to group differentiation.

Overall, separation between ECs and ART-suppressed non-controllers was greatest when clinical covariates were considered together with the top 15 differentially expressed proteins. The area under the receiver operating characteristic curve (auROC) was 0.822 (95% CI 0.73–0.91), where 1.0 indicates perfect separation and 0.5 indicates chance-level performance (**Figure 7A**). The area under the precision-recall curve (auPR) was 0.856 (95% CI 0.75–0.94), compared with a baseline of 0.626. Features contributing most strongly to differentiation of ECs included higher CD4+ T-cell counts, lower levels of intact provirus, and lower levels of the chemokines CXCL12, CCL23, and CCL24. ECs were also characterized by lower levels of angiogenesis/extracellular matrix–related proteins (VEGF-A, SMOC2, AGRN) and gut mucosal/intestinal epithelial–associated proteins (SPINK4, LGALS4, REG4) (**Figure 7B**). These findings characterize the combination of clinical and proteomic features associated with the EC phenotype in this cohort and should not be interpreted as establishing an independently validated predictive or classification signature.

**Figure 7.**
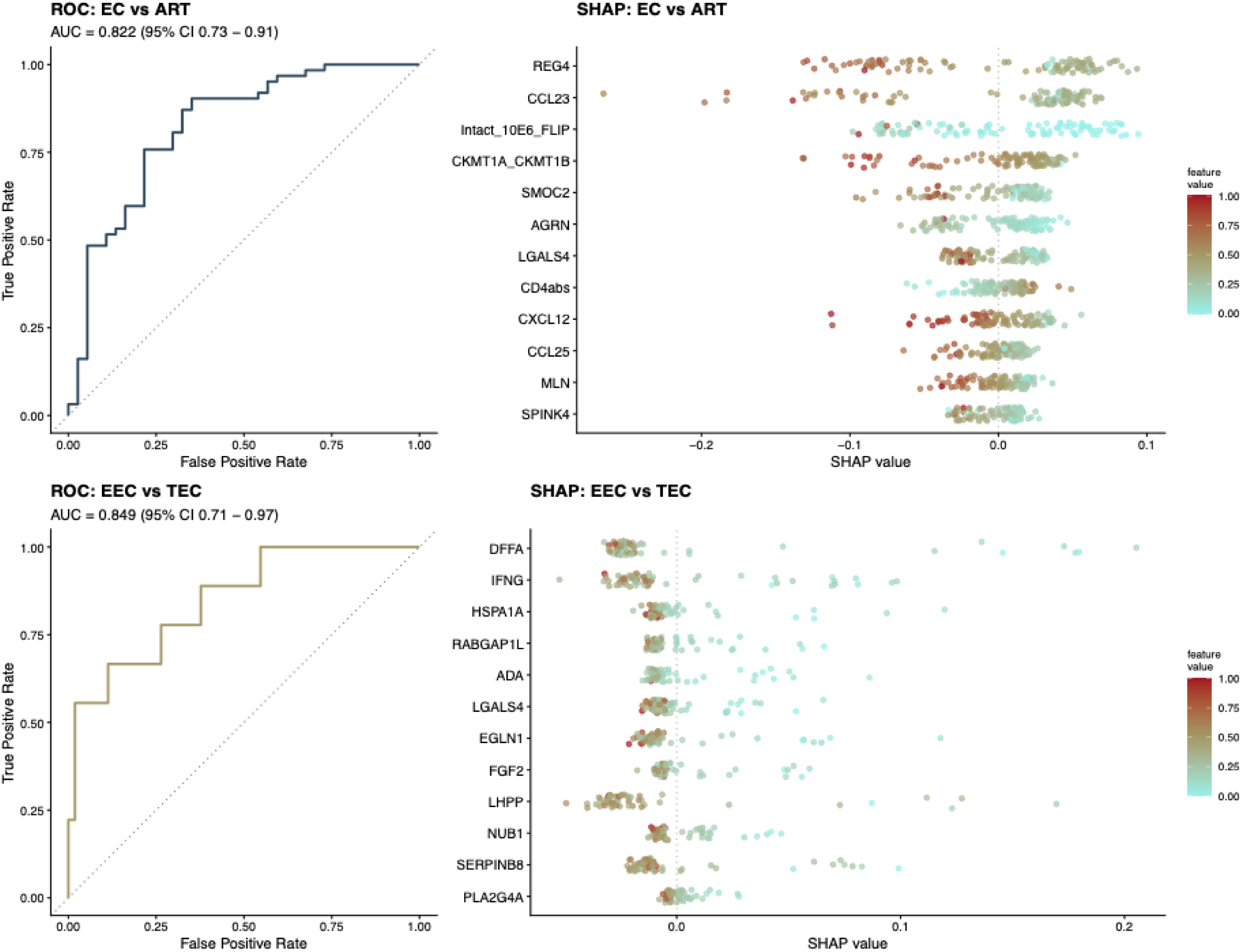
Machine-learning characterization of clinical and proteomic features associated with elite controller phenotypes. **A–B.** A nested cross-validation Random Forest model with SHapley Additive exPlanations (SHAP) was used to distinguish ART-naïve elite controllers (ECs; n = 62) from ART-suppressed non-controllers (n = 37) using the top 15 differentially expressed Olink proteins (selected by auROC over {10, 15, 20}; see Methods), absolute CD4 count, and intact proviral DNA. A. Receiver operating characteristic (ROC) curve showing separation between ECs and ART-suppressed non-controllers (auROC = 0.822, 95% CI 0.73–0.91). B. SHAP summary plot highlighting features contributing most strongly to differentiation of ECs, including higher CD4 counts, lower intact proviral DNA, lower levels of chemokines (e.g., CXCL12, CCL23, CCL24), and lower levels of angiogenesis/extracellular matrix–related proteins and gut mucosal/intestinal epithelial–associated proteins. **C–D.** The same nested cross-validation RF–SHAP workflow was applied to characterize differences between ECs with PWoH-like proteomic profiles (n = 9) and typical ECs (n = 53) using clinical covariates plus the 20 proteins most strongly distinguishing the predefined subgroups (selected by auROC over {10, 15, 20}; see Methods). **C.** ROC curve showing within-cohort separation of the predefined EC subgroups (auROC = 0.849, 95% CI 0.71–0.97). **D.** SHAP summary plot highlighting features contributing to subgroup differentiation, including lower inflammatory, apoptosis, and stress-related markers (e.g., DFFA, IFNG, HSPA1A, ADA, PLA2G4A), as well as lower LGALS4 expression, among ECs with PWoH-like proteomic profiles.

### Machine-learning analysis characterizes features associated with PWoH-like proteomic profiles among ECs

We next applied the same approach to characterize the features that most strongly distinguished ECs with PWoH-like proteomic profiles from typical ECs. Because the EC subgroups were defined using the same proteomic measurements subsequently included in these analyses, this analysis was intended as hypothesis-generating characterization of the features associated with the observed proteomic states rather than as independent classification or prediction. Separation was greatest when clinical covariates were considered together with the 20 proteins most strongly distinguishing the predefined subgroups (auROC = 0.849 [95% CI 0.71–0.97], auPR = 0.638 [95% CI 0.30–0.90]; baseline 0.145; **Figure 7C**).

Features contributing most strongly to differentiation of ECs with PWoH-like proteomic profiles included higher CD4+ T-cell counts and lower levels of inflammatory markers, including IFNG, ADA, PLA2G4A, and FGF2. HSPA1A also contributed to distinguishing ECs with PWoH-like proteomic profiles from typical ECs (Figure 7D). Consistent with the EC versus ART-suppressed comparison (**Figure 7B**), lower expression of the gut mucosal/intestinal epithelial–associated protein LGALS4 (galectin-4) was characteristic of ECs with PWoH-like proteomic profiles compared with typical ECs.

We further explored whether individual proteins could characterize the distinction between ECs with PWoH-like proteomic profiles and typical ECs. To complement the differential expression and multivariable analyses, we used a simple one-feature approach (OneRule/OneR), which evaluates each protein individually and identifies the expression threshold that best separates the predefined EC subgroups within the dataset (**Supplementary Figure 2**). This exploratory approach can identify proteins that are not among the most differentially expressed based on average group differences but nonetheless strongly characterize the observed subgroup separation. HCLS1 provided the strongest single-feature separation, with lower HCLS1 expression characteristic of ECs with PWoH-like proteomic profiles. An HCLS1 NPX threshold of < –0.7819 provided the greatest separation of the two predefined EC subgroups in this dataset (**Supplementary Figure 2**). These findings identify reduced HCLS1 as a candidate protein associated with the PWoH-like proteomic state.

## DISCUSSION

There is continued interest in understanding ECs as a model for functional cure of HIV infection, but outcomes among ECs are heterogeneous. Identifying immunologic and proteomic correlates of control could clarify whether some ECs reach a state resembling a cure. Using plasma proteomics, we performed differential expression and machine learning analyses and found that ART-naïve ECs exhibited lower levels of inflammation than ART-suppressed non-controllers but higher levels of inflammation than people without HIV. We further identified a subset of ECs that resembled people without HIV proteomically. Importantly, these ECs with PWoH-like proteomic profiles were characterized by numerically lower intact and total proviral reservoirs and a high frequency of protective HLA alleles. Within this phenotype lower levels of inflammatory proteins, notably IL-1β, and reduced HCLS1 expression were prominent features of this subgroup, alongside lower levels of select select gut mucosal/intestinal epithelial–associated proteins in plasma compared with typical ECs. Together, our data add additional insight to evidence that a subset of elite controllers may achieve a near-cure phenotype [16–23].

Consistent with previous studies, we observed that compared to non-controllers on ART, ECs had lower intact proviral frequencies [19, 25, 26] and higher absolute CD4+ T-cell counts [9, 27]. Proteomic analyses revealed that markers of immune activation (CXCL12, CCL23, CCL24), angiogenesis/ECM remodeling (VEGF-A, SMOC2, AGRN), and gut mucosa (SPINK4, LGALS4, REG4) were downregulated in ECs relative to ART-suppressed non-controllers. Results of prior studies are mixed as to whether ECs have higher or lower levels of inflammation relative to ART-suppressed non-controllers, likely reflecting biological heterogeneity within ECs as well as differences in cohort composition, cohort scale, and analytic approaches [8–12, 16–23, 27–31]. The direction of group-level differences may depend in part on the composition of the EC population sampled, particularly when relatively small numbers of individuals are available for study. Given the substantial heterogeneity we observed among 62 ECs, differential representation of low- and high-inflammatory EC phenotypes across cohorts could plausibly contribute to apparently discordant comparisons with ART-suppressed individuals [8–12, 16–23, 27–31]. However, it is notable from our study that ECs, when analyzed as a group, still exhibited upregulation of multiple inflammatory mediators relative to PWoH, including pathways linked to cytokine signaling, immune activation, cellular stress, and apoptosis. This suggests that even the relatively low reservoir state of ECs can be associated with inflammation and that a clinical state resembling virologically-defined elite control may not be the optimal goal for a curative intervention [32].

Our clustering analyses revealed substantial heterogeneity among ECs, with a subset showing proteomic profiles closely resembling those of people without HIV, consistent with prior transcriptional studies [33]. Differential expression analysis identified three proteins that seemed to most consistently define ECs with PWoH-like proteomic profiles: IL-1β, SCRN1, and BID. Notably, IL-1β is a central upstream mediator of inflammation and has clinical relevance: in the CANTOS trial, pharmacologic IL-1β blockade with canakinumab reduced systemic inflammatory markers and lowered the risk of recurrent atherosclerotic cardiovascular events [34, 35]. In HIV, IL-1β-driven pyroptosis contributes to CD4 T-cell loss [36], and downstream mediators such as RANTES have been associated with disease progression in ECs [37]. SCRN1, which has been associated with chronic inflammatory states and HIV disease progression, and BID, a pro-apoptotic regulator, were also consistently reduced, consistent with a proteomic profile characterized by lower inflammatory and apoptosis-related markers [38, 39]. Importantly, these findings should be interpreted in the context of a longitudinal proteomic study of elite and viremic controllers, which indicated that systemic inflammatory mediators, including IL-1β, do not predict subsequent loss of viral control over extended follow-up [40]. However, a distinct PWoH-like inflammatory proteomic state may nevertheless characterize a rare subset of ECs.

Additionally, ECs with PWoH-like proteomic profiles exhibited lower expression of gut injury-associated proteins, including LGALS4 (galectin-4) [41, 42], raising the hypothesis that differences in mucosal integrity may contribute to the observed systemic inflammatory profile, as suggested by prior studies linking galectins and mucosal barrier disruption to inflammasome activation in HIV and SIV infection [36, 43, 44]. Notably, most ECs with PWoH-like proteomic profiles carried at least one protective HLA allele, suggesting that effective CD8⁺ T cell–mediated viral control may contribute to this low-inflammatory state, rather than inflammation being suppressed solely through innate immune pathways [21, 45–51]. The high frequency of protective HLA alleles, together with numerically lower total reservoir frequencies by FLIP-seq, is also potentially consistent with early control of HIV viremia driven by strong T cell responses in these nine ECs with PWoH-like proteomic profiles.

Interestingly, lower expression of HCLS1 (hematopoietic cell-specific Lyn substrate 1) provided the strongest single-feature separation of ECs with PWoH-like proteomic profiles compared to typical ECs. HCLS1 is an actin-binding protein expressed in hematopoietic cells and has multiple roles in immune signaling [52, 53]. It has also been implicated in the dissemination of latent CMV and is upregulated in the rectosigmoid tissue of HIV non-controllers relative to ECs [52–54]. Therefore, we speculate that reduced HCLS1 expression in ECs with PWoH-like proteomic profiles may be associated with dampened inflammatory signaling. Future studies may evaluate whether decreased HCLS1 expression limits inflammatory signaling and thereby influences reservoir maintenance or expansion.

This study has several limitations. First, the cross-sectional design precludes inference about causality or the temporal relationship between inflammation, reservoir size, and virologic control. Second, the small number of ECs with PWoH-like proteomic profiles limits statistical power and resulted in substantial uncertainty in exploratory model performance (auROC 0.849, 95% CI 0.71–0.97). Third, the EC subgroups were defined by clustering of the same proteomic measurements subsequently used to characterize them. Thus, differential-expression, machine-learning, and single-feature analyses involving these subgroups are inherently circular and should be interpreted as descriptive, hypothesis-generating analyses rather than independent classification or prediction. Fourth, the Olink Inflammation I Explore Panel captures a pre-selected subset of proteins and therefore does not represent the full proteome [55]. Fifth, the rarity of this phenotype makes fully independent discovery and validation cohorts difficult to assemble; nonetheless, confirmation in larger independent cohorts will be important to determine whether these proteomic features reproducibly define a PWoH-like state and whether they relate to clinically meaningful outcomes. Sixth, we were unable to assess clinical outcomes related to inflammation. Finally, the predominance of non-Hispanic White male participants limits generalizability and highlights the need for more diverse and representative study populations.

In summary, we identified a distinct proteomic profile in a subset of ECs whose inflammatory signatures closely resemble those of people without HIV. These findings extend virologic characterization of elite control by identifying specific immune and signaling pathways associated with a PWoH-like inflammatory state and provide candidate pathways for further investigation in studies of HIV remission and cure.

## METHODS

### Study Participants and Ethics Statement

This study involved human participants and was conducted in accordance with the principles of the Declaration of Helsinki. All participants were enrolled under protocols approved by the Institutional Review Boards (IRBs) of the University of California, San Francisco (UCSF), Massachusetts General Hospital, and Brigham and Women’s Hospital. Informed consent was obtained from all participants prior to inclusion in the study. All samples and associated clinical data were de-identified prior to analysis to protect participant confidentiality. The use of stored biospecimens and clinical data for proteomic and genomic analyses was approved by the respective IRBs.

We studied ART-naïve aviremic ECs (n = 62) and ART-suppressed non-controllers (n = 37) from a previously described cohort [19]. Individuals without HIV (n = 50) from the UCSF SCOPE cohort served as comparators. Informed consent was obtained from all human research participants.

### Clinical Data and HIV Reservoir Measurements

Demographic and clinical data were obtained from clinical repositories, and absolute CD4 counts and CD4:CD8 ratios were measured using standard assays. Intact HIV DNA in PBMCs was quantified using FLIP-Seq, a near–full-length single-genome sequencing method for reservoir assessment [56]. Detailed FLIP-Seq results for this cohort have been reported previously [19]. Absolute CD4+ T-cell counts, CD4:CD8 ratios, time since HIV diagnosis, and intact provirus levels were compared between groups using linear regression adjusted for age and gender [57]. Outliers were identified using the 1.5× IQR method, and normality was assessed with Shapiro-Wilk or Lilliefors tests in R. Normally distributed data without outliers were analyzed using t-tests; otherwise, comparisons used Mann-Whitney U tests in R.

### Olink Proteomics

We measured 384 plasma proteins using the Olink Inflammation Explore Panel [55]. Normalized Protein eXpression (NPX) values were processed according to the manufacturer’s Intensity Normalization protocol, assuming no median differences across sample-randomized plates. For each assay, we calculated sample and plate medians, derived a plate-specific median, subtracted this value from each measurement on that plate, and added the overall assay median to generate normalized NPX values.

### Differential NPX and Enrichment Analysis

Differential NPX values were analyzed using the OlinkAnalyze R package. Quality control and data distribution were assessed with ‘olink_qc_plot,’ ‘olink_dist_plot,’ and PCA/UMAP visualizations. After filtering to the comparator groups, differential protein expression was assessed using the parametric ‘olink_ttest’ function. Effect estimates and Benjamini–Hochberg–adjusted p-values were used to generate volcano plots (ggplot2) and for downstream analyses.

Gene Ontology (GO) enrichment analysis was performed across the three GO domains, Biological Process (BP), Molecular Function (MF), and Cellular Component (CC), using the enrichGO function, followed by redundancy reduction with simplify. Kyoto Encyclopedia of Genes and Genomes (KEGG) pathway enrichment was conducted using enrichKEGG, with statistical significance assessed using Benjamini–Hochberg–adjusted p-values.

### Heatmap Clustering

ANOVA was used to evaluate model fit for each protein using two models: (1) NPX ∼ Gender and (2) NPX ∼ Cohort + Gender. Model comparison tested whether adding ‘Cohort’ explained additional variance (FDR < 0.05). Proteins meeting this criterion were flagged as cohort-associated.

These cohort-associated proteins were visualized in a heatmap comparing ECs, ART-suppressed non-controllers, and PWoH. Proteins were clustered by NPX values using k-means with the optimal cluster number determined by the hopach package, and samples were hierarchically clustered using pheatmap.

A second, descriptive heatmap examined ECs and PWoH only. Because patterns could reflect age or gender, we also performed pairwise regression (NPX ∼ Cohort + Gender + Age) to estimate cohort effects adjusted for these covariates [57]. This third heatmap allows for a more focused look at differences between ECs and individuals without HIV, adjusted for potential confounders.

From the heatmaps, ECs clustering within the predominant PWoH cluster in at least one of the three analyses were designated ECs with PWoH-like proteomic profiles (n = 9); the remaining ECs were designated typical ECs (n = 53).

### Machine Learning

To understand the discriminatory value and individual contributions of proteins beyond univariate analysis, we trained Random Forest (RF) models using ranger for exploratory characterization. Because subgroup labels were derived from clustering of the same proteomic measurements subsequently used as predictors, cross-validation cannot provide an independent estimate of classification performance for this comparison. Accordingly, these analyses were treated as descriptive feature-characterization analyses rather than validated predictive models.

Each RF comparison included the top 10, 15, and 20 proteins ranked by t-test within each cross validation training fold, and combined with clinical covariates (HIV reservoir frequency and absolute CD4 counts). For each comparison (EC vs ART; ECs with PWoH-like proteomic profiles vs typical EC), the protein-set size (10, 15, or 20) with the highest pooled auROC was reported as exploratory.

Each RF model (n=1000 trees) was applied with nested cross-validation to characterize how selected proteins discriminate when considered collectively. The outer loop consisted of 10-fold cross-validation (CV) repeated 10 times to estimate model performance, while the inner loop (10-fold CV) selected hyperparameters (mtry, min.node.size, max.depth, sample.fraction). For training of each outer fold, the top differential proteins were re-ranked by t-test on that fold’s training samples only. All preprocessing was performed using training-set statistics only within each cross-validation fold [58]. Stratified 10-fold cross-validation was repeated 10 times. Each participant was held out once per repeat, yielding 10 out-of-fold predicted probabilities per participant; these probabilities were averaged to obtain a single participant-level prediction for calculation of auROC and auPR. 95% CIs were generated using 1,000 bootstrap replicates.

Feature importance was determined using SHapley Additive exPlanations (SHAP) values combined across the outer folds. The mean |SHAP| value was used for feature importance. Feature stability was determined by the number of outer folds in which each feature was selected into the top N by the within fold t-test ranking. These models characterize discriminatory patterns rather than provide generalizable prediction. For visualization, feature values were min-max normalized within each protein to a 0–1 scale for the SHAP color gradient; SHAP values themselves were not normalized.

For additional simplified comparison of ECs with PWoH-like proteomic profiles and typical ECs, we further applied OneR in the Waikato Environment for Knowledge Analysis (WEKA). The OneR model included all proteins in the panel and the above clinical covariates. OneR was selected because it generates highly interpretable, single-rule models, allowing us to identify the individual protein or clinical variable providing the strongest single-feature separation between the predefined EC subgroups.

## Acknowledgements

Graphical Abstract Created with BioRender.com. We are grateful to the study participants.

## Data Availability

Analysis code is available at: https://github.com/davidpmaison/falcinelli_peluso_elitecontrollers.

---

**Supplementary Figure 1.**
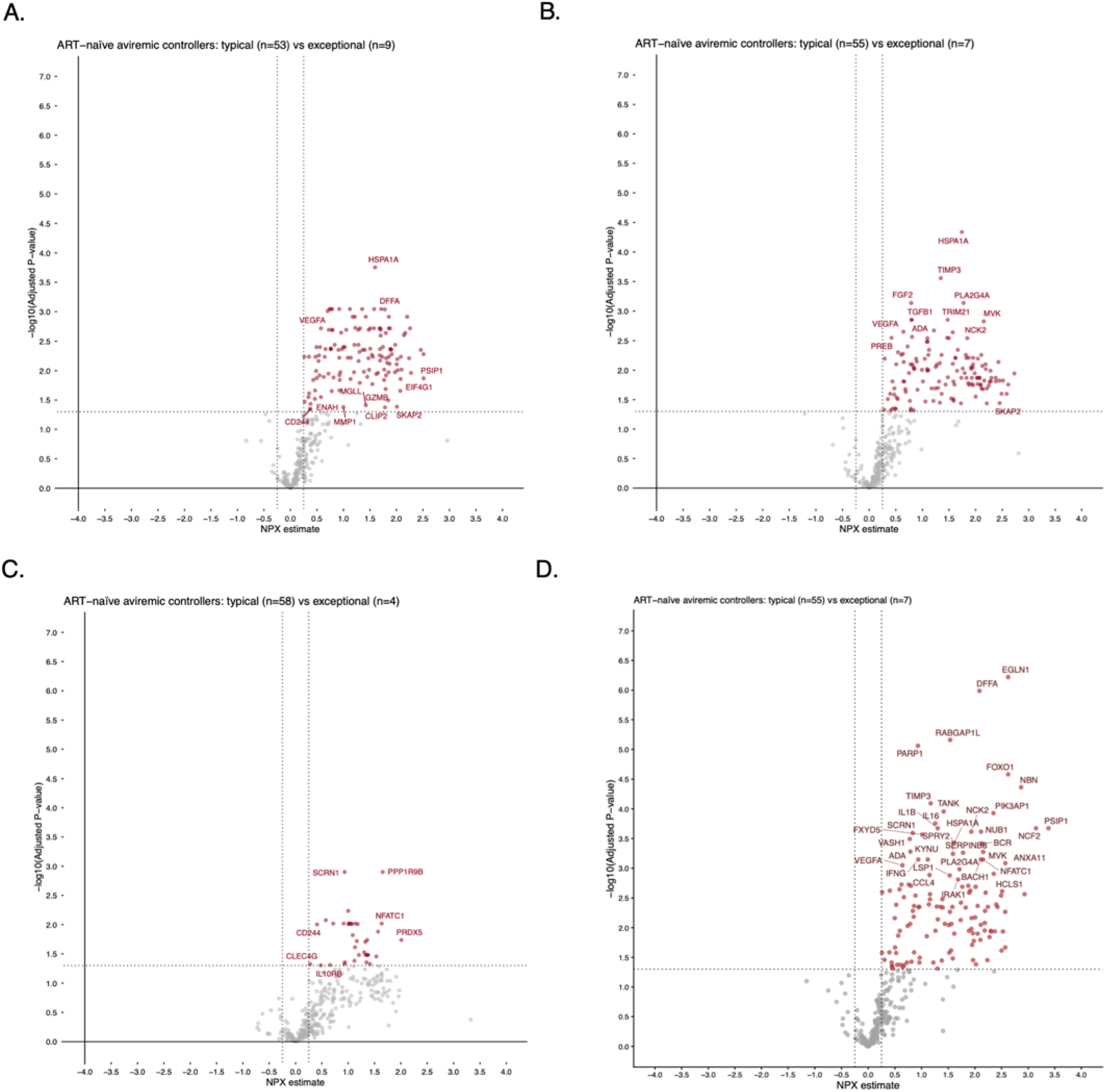
IL-1β, SCRN1, BID proteins are consistently differentially expressed in ECs with PWoH-like proteomic profiles. To identify protein expression patterns that distinguish ECs with PWoH-like proteomic profiles from typical ECs, we compared results across three complementary clustering and differential expression approaches. IL-1β, SCRN1, and BID were repeatedly among the top 15 most significantly differentially expressed proteins (adjusted p-value), with all upregulated proteins enriched in typical ECs. **A.** Differential expression analysis comparing ECs with PWoH-like proteomic profiles (n = 9) versus typical ECs (n = 53). **B.** Differential expression results for ECs with PWoH-like proteomic profiles identified through clustering based on significant cohort-associated proteins derived from comparisons among ECs, ART-suppressed non-controllers, and people without HIV. **C.** Differential expression results for ECs with PWoH-like proteomic profiles identified by unsupervised clustering with PWoH using all measured proteins. **D.** Differential expression results for ECs with PWoH-like proteomic profiles identified by unsupervised clustering with PWoH individuals using all measured proteins, adjusted for age and gender.

**Supplementary Figure 2.**
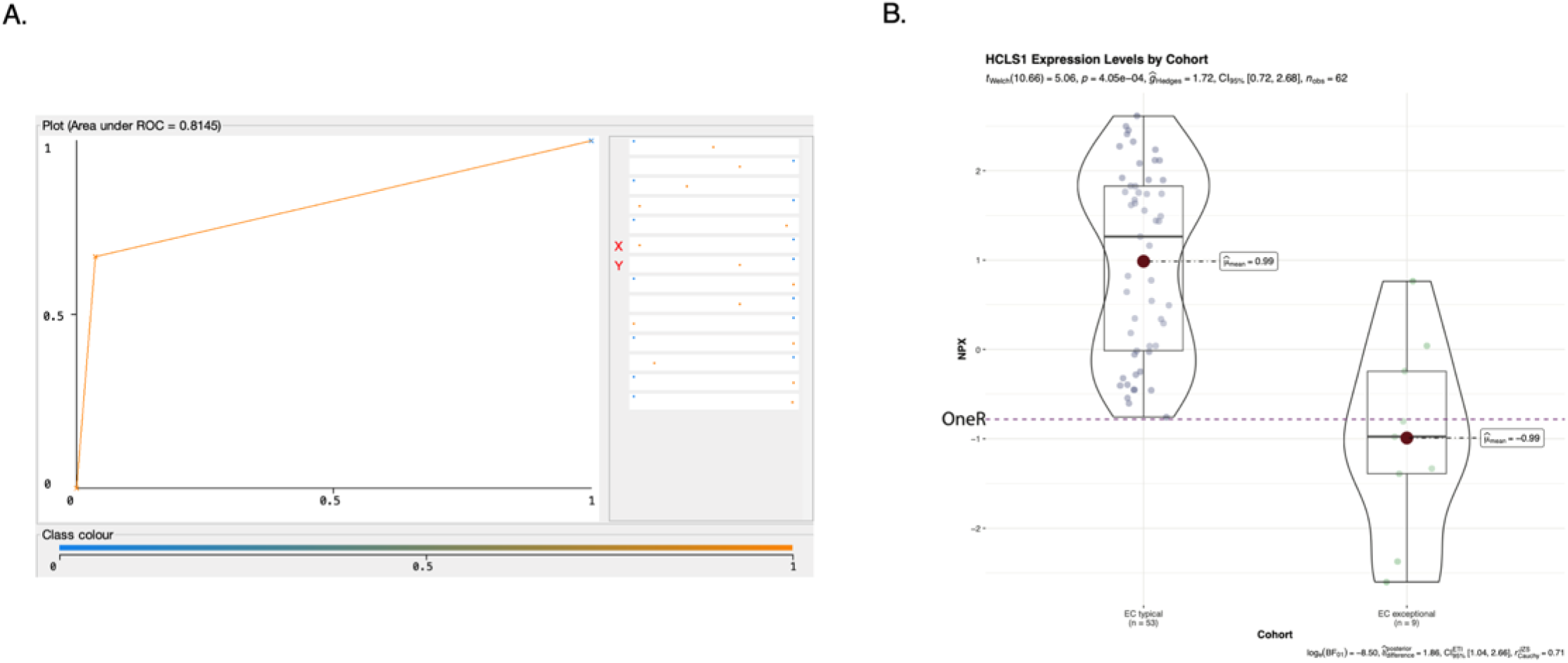
OneR analysis identifies HCLS1 as the single feature providing the strongest separation of predefined EC subgroups. To explore whether an individual feature could characterize differences between ECs with PWoH-like proteomic profiles (n = 9) and typical ECs (n = 53), we applied the OneRule (OneR) algorithm to evaluate each feature individually. HCLS1 provided the strongest single-feature separation in this dataset, with lower HCLS1 expression among ECs with PWoH-like proteomic profiles. OneR identified an NPX threshold of –0.7819 as providing the greatest within-dataset separation between the predefined subgroups. **A.** ROC curve describing within-dataset separation based on HCLS1 (auROC = 0.8145). **B.** Violin/box plot of HCLS1 NPX values in typical ECs and ECs with PWoH-like proteomic profiles. The dashed horizontal line indicates the exploratory OneR threshold (NPX = –0.7819).

**Supplementary Table 1.** Protective Class I HLA alleles by cohort.

| Cohort | HLA<br>Class1A | HLA<br>Class1A2 | HLA<br>Class1B | HLA<br>Class1B2 | HLA<br>Class1CW | HLA<br>Class1CW<br>2 | Protectiv<br>e Allele | 2nd<br>Protectiv<br>e Allele |
| --- | --- | --- | --- | --- | --- | --- | --- | --- |
| EC resembling PWoH | 2:01 | 11:01 | 27:05:00 | 40:01:00 | 1:02 | 3:04 | 27:05:00 |  |
| EC resembling PWoH | 3:01 | 31:01:00 | 27:05:00 | 57:01:00 | 3:07 | 6:02 | 57:01:00 |  |
| EC resembling PWoH | 2:01 | 36:01:00 | 40:02:00 | 57:01:00 | 3:04 | 6:02 | 57:01:00 |  |
| EC resembling PWoH | 1:01 | 24:02:00 | 38:01:00 | 57:01:00 | 6:02 | 12:03 | 57:01:00 |  |
| EC resembling PWoH | 2:02 | 2:11 | 45:01:00 | 57:03:00 | 7:01 | 16:01 | 57:03:00 |  |
| EC resembling PWoH | 31:01:00 | 36:01:00 | 57:03:00 | 58:02:00 | 6:02 | 7:01 | 57:03:00 |  |
| EC resembling PWoH | 23:01 | 25:01:00 | 44:03:00 | 58:01:00 | 4:01 | 7:01 | 58:01:00 |  |
| EC resembling PWoH | 11:01 | 32:01:00 | 35:01:00 | 50:01:00 | 4:01 | 6:02 |  |  |
| EC resembling PWoH | 68:01:00 | 68:02:00 | 15:10 | 51:02:00 | 3:04 | 16:01 |  |  |
| ART | 2:01 | 11:01 | 27:05:00 | 44:02:00 | 1:02 | 5:01 | 27:05:00 |  |
| ART | 1:01 | 3:01 | 27:05:00 | 39:10:00 | 2:02 | 12:03 | 27:05:00 |  |
| ART | 1:01 | 2:01 | 44:02:00 | 57:01:00 | 6:02 | 12:03 | 57:01:00 |  |
| ART | 1:01 | 25:01:00 | 8:01 | 57:01:00 | 6:02 | 7:01 | 57:01:00 |  |
| ART | 1:01 | 3:01 | 7:02 | 57:01:00 | 6:02 | 7:02 | 57:01:00 |  |
| ART | 3:01 | 36:01:00 | 35:02:00 | 57:03:00 | 4:01 | 7:01 | 57:03:00 |  |
| ART | 30:02:00 | 66:01:00 | 15:10 | 57:03:00 | 3:04 | 18 | 57:03:00 |  |
| ART | 1:01 | 31:01:00 | 40:01:00 | 58:01:00 | 3:02 | 3:04 | 58:01:00 |  |
| ART | 2:01 | 3:01 | 7:02 | 8:01 | 7:01 | 7:02 |  |  |
| ART | 33:01:00 | 74:01:00 | 14:02 | 42:01:00 | 8:02 | 17 |  |  |
| ART | 3:01 | 24:02:00 | 7:02 | 39:06:00 | 7:02 | 7:02 |  |  |
| ART | 2:01 | 11:01 | 35:03:00 | 40:01:00 | 3:04 | 4:01 |  |  |
| ART | 29:01:00 | 32:01:00 | 7:02 | 40:01:00 | 3:04 | 7:02 |  |  |
| ART | 30:01:00 | 30:02:00 | 15:10 | 35:01:00 | 3:04 | 4:01 |  |  |
| ART | 2:01 | 3:01 | 14:02 | 44:02:00 | 5:01 | 8:02 |  |  |
| ART | 3:01 | 30:01:00 | 7:02 | 35:01:00 | 4:01 | 7:02 |  |  |
| ART | 11:01 | 24:02:00 | 7:05 | 13:01 | 3:04 | 7:02 |  |  |
| ART | 2:01 | 2:01 | 15:01 | 40:01:00 | 3:03 | 3:04 |  |  |
| ART | 11:01 | 68:01:00 | 15:01 | 18:03 | 3:03 | 7:01 |  |  |
| ART | 1:01 | 2:01 | 7:02 | 15:01 | 3:04 | 7:02 |  |  |
| ART | 3:01 | 30:02:00 | 7:02 | 18:01 | 5:01 | 7:02 |  |  |
| ART | 23:01 | 80:01:00 | 8:01 | 52:01:00 | 7:01 | 16:01 |  |  |
| ART | 2:01 | 2:01 | 15:01 | 44:02:00 | 3:04 | 12:03 |  |  |
| ART | 29:02:01 | 310102 | 3512 | 440301 | 401 | 160101 |  |  |
| ART | 23:01 | 29:02:00 | 15:01 | 53:01:00 | 3:03 | 4:01 |  |  |
| ART | 1:01 | 31:01:00 | 8:01 | 39:01:00 | 7:01 | 7:02 |  |  |
| ART | 30:02:00 | 74 | 7:05 | 40:16:00 | 8:02 | 15:05 |  |  |
| ART | 2:01 | 2:01 | 7:02 | 44:02:00 | 5:01 | 7:02 |  |  |
| ART | 1:01 | 24:02:00 | 15:17 | 39:06:00 | 7:01 | 7:02 |  |  |
| ART | 3:01 | 24:02:00 | 14:02 | 55:01:00 | 3:04 | 8:02 |  |  |
| ART | 2:17 | 29:02:00 | 44:03:00 | 51:01:00 | 15:02 | 16:01 |  |  |
| ART | 2:01 | 32:01:00 | 14:01 | 44:02:00 | 5:01 | 8:02 |  |  |
| ART | 2:02 | 30:01:00 | 7:02 | 35:01:00 | 7:05 | 7:05 |  |  |
| ART | 1:01 | 24:02:00 | 8:01 | 44:03:00 | 7:01 | 16:01 |  |  |
| ART | 3:01 | 30:02:00 | 7:02 | 18:01 | 5:01 | 7:02 |  |  |
| ART | 2:01 | 11:01 | 40:01:00 | 55:01:00 | 3:03 | 3:04 |  |  |
| ART | 2:01 | 3:01 | 38:01:00 | 52:01:00 | 12:03 | 16:01 |  |  |
| Typical EC | 1:01 | 11:01 | 27:05:00 | 35:03:00 | 2:02 | 4:01 | 27:05:00 |  |
| Typical EC | 2:06 | 24:02:00 | 1/1/00<br>3:05 | 40:06:00 | 2:02 | 8:01 | 27:05:00 |  |
| Typical EC | 2:01 | 11:01 | 15:18 | 27:05:00 | 1:02 | 7:04 | 27:05:00 |  |
| Typical EC | 2:01 | 3:01 | 15:01 | 27:05:00 | 1:02 | 3:04 | 27:05:00 |  |
| Typical EC | 2:01 | 11:01 | 14:02 | 51:01:00 | 8:02 | 14:02 | 51:01:00 |  |
| Typical EC | 3:01 | 26:01:00 | 44:02:00 | 51:01:00 | 5:01 | 15:02 | 51:01:00 |  |
| Typical EC | 1:01 | 2:01 | 51:01:00 | 57:01:00 | 6:02 | 14:02 | 57:01:00 |  |
| Typical EC | 1:01 | 3:01 | 41:02:00 | 57:01:00 | 6:02 | 12:02 | 57:01:00 |  |
| Typical EC | 2:01 | 23:01 | 49:01:00 | 57:01:00 | 6:02 | 7:01 | 57:01:00 |  |
| Typical EC | 2:01 | 2:01 | 27:05:00 | 57:01:00 | 1:02 | 6:02 | 57:01:00 | 27:05:00 |
| Typical EC | 11:01 | 31:01:00 | 38:01:00 | 57:01:00 | 6:02 | 12:03 | 57:01:00 |  |
| Typical EC | 24:02:00 | 24:02:00 | 14:02 | 57:01:00 | 6:02 | 8:02 | 57:01:00 |  |
| Typical EC | 1:01 | 34:02:00 | 8:01 | 57:01:00 | 6:02 | 7:01 | 57:01:00 |  |
| Typical EC | 1:01 | 30:01:00 | 13:02 | 57:01:00 | 6:02 | 6:02 | 57:01:00 |  |
| Typical EC | 1:01 | 68:01:00 | 55:01:00 | 57:01:00 | 3:03 | 6:02 | 57:01:00 |  |
| Typical EC | 3:01 | 26:01:00 | 14:01 | 57:01:00 | 6:02 | 8:02 | 57:01:00 |  |
| Typical EC | 2:01 | 30:01:00 | 13:02 | 57:01:00 | 6:02 | 6:02 | 57:01:00 |  |
| Typical EC | 1:01 | 3:01 | 7:02 | 57:01:00 | 6:02 | 7:02 | 57:01:00 |  |
| Typical EC | 2:05 | 30:01:00 | 42:01:00 | 57:03:00 | 17 | 18 | 57:03:00 |  |
| Typical EC | 30:02:00 | 33:03:00 | 15:16 | 57:03:00 | 7:01 | 14:02 | 57:03:00 |  |
| Typical EC | 2:02 | 74 | 49:01:00 | 57:03:00 | 7:01 | 7:01 | 57:03:00 |  |
| Typical EC | 32:01:00 | 68:02:00 | 39:06:00 | 57:03:00 | 7:01 | 7:02 | 57:03:00 |  |
| Typical EC | 30:01:00 | 33:03:00 | 42:01:00 | 57:03:00 | 17 | 18 | 57:03:00 |  |
| Typical EC | 1:01 | 31:01:00 | 53:01:00 | 57:03:00 | 12:02 | 12:03 | 57:03:00 |  |
| Typical EC | 2:01 | 30:02:00 | 44:02:00 | 57:03:00 | 5:01 | 18 | 57:03:00 |  |
| Typical EC | 30:01:00 | 74 | 35:01:00 | 57:03:00 | 16:01 | 18 | 57:03:00 |  |
| Typical EC | 3:01 | 33:03:00 | 57:02:00 | 58:01:00 | 7:01 | 18 | 58:01:00 |  |
| Typical EC | 1:01 | 29:02:00 | 44:03:00 | 58:01:00 | 6:02 | 16:01 | 58:01:00 |  |
| Typical EC | 2:01 | 25:01:00 | 44:02:00 | 58:01:00 | 5:01 | 7:01 | 58:01:00 |  |
| Typical EC | 1:01 | 11:01 | 35:01:00 | 58:01:00 | 4:01 | 7:01 | 58:01:00 |  |
| Typical EC | 2:01 | 24:02:00 | 27:05:00 | 58:01:00 | 1:02 | 6:02 | 58:01:00 | 27:05:00 |
| Typical EC | 2:02 | 24:02:00 | 58:01:00 | 81:01:00 | 3:04 | 5:01 | 58:01:00 |  |
| Typical EC | 1:01 | 11:01 | 8:01 | 18:01 | 7:01 | 7:01 |  |  |
| Typical EC | 3:01 | 31:01:00 | 14:01 | 35:01:00 | 4:01 | 8:02 |  |  |
| Typical EC | 1:03 | 32:01:00 | 35:08:00 | 73:01:00 | 4:01 | 15:05 |  |  |
| Typical EC | 2:01 | 3:01 | 51:01:00 | 51:01:00 | 14:02 | 15:02 |  |  |
| Typical EC | 24:02:00 | 25:01:00 | 39:11:00 | 44:03:00 | 7:02 | 16:01 |  |  |
| Typical EC | 68:01:00 | 69:01:00 | 35:03:00 | 39:01:00 | 4:01 | 12:03 |  |  |
| Typical EC | 3:01 | 32:01:00 | 14:02 | 40:01:00 | 3:04 | 8:02 |  |  |
| Typical EC | 11:01 | 24:02:00 | 15:01 | 35:01:00 | 3:03 | 4:01 |  |  |
| Typical EC | 2:01 | 3:01 | 7:02 | 14:02 | 7:02 | 8:02 |  |  |
| Typical EC | 23:01 | 34:02:00 | 52:01:00 | 81:01:00 | 8:04 | 16:01 |  |  |
| Typical EC | 3:01 | 11:01 | 35:01:00 | 51:01:00 | 4:01 | 15:02 |  |  |
| Typical EC | 2:01 | 3:01 | 15:01 | 51:01:00 | 3:03 | 15:02 |  |  |
| Typical EC | 2:01 | 3:01 | 7:02 | 52:01:00 | 7:02 | 12:02 |  |  |
| Typical EC | 24:02:00 | 30:02:00 | 18:01 | 53:01:00 | 4:01 | 12:03 |  |  |
| Typical EC | 24:02:00 | 31:01:00 | 40:01:00 | 44:02:00 | 3:04 | 5:01 |  |  |
| Typical EC | 1:01 | 74 | 37:01:00 | 81:01:00 | 6:02 | 18 |  |  |
| Typical EC | 1:01 | 1:01 | 8:01 | 35:02:00 | 6:02 | 7:01 |  |  |
| Typical EC | 3:01 | 29:02:00 | 15:01 | 45:01:00 | 3:03 | 6:02 |  |  |
| Typical EC | 3:01 | 66:01:00 | 41:02:00 | 49:01:00 | 4:01 | 17 |  |  |
| Typical EC | 2:01 | 24:02:00 | 44:02:00 | 57:03:00 | 5:01 | 7:01 |  |  |
| Typical EC | 2:01 | 31:01:00 | 15:01 | 44:02:00 | 3:03 | 5:01 |  |  |

